# Identifying low risk insecticides that can enhance food production without increasing mortality of biocontrol agents for human schistosomiasis

**DOI:** 10.1101/2021.01.05.425425

**Authors:** Christopher J E Haggerty, Bryan K. Delius, Nicolas Jouanard, Pape D Ndao, Giulio A De Leo, Andrea J Lund, David Lopez-Carr, Justin V Remais, Gilles Riveau, Susanne H Sokolow, Jason R Rohr

## Abstract

Use of agrochemicals, including insecticides, is vital to food production and predicted to increase 2-5 fold by 2050. Previous studies have shown a positive association between agriculture and the human infectious disease schistosomiasis, which is problematic as this parasitic disease infects approximately 250 million people worldwide. Certain insecticides might runoff fields and be highly toxic to invertebrates, such as prawns in the genus *Macrobrachium*, that are biocontrol agents for snails that transmit the parasites causing schistosomiasis. We used a laboratory dose-response experiment and an observational field study to determine the relative toxicities of three pyrethroid (esfenvalerate, λ-cyhalothrin, and permethrin) and three organophosphate (chlorpyrifos, malathion, and terbufos) insecticides to *Macrobrachium* prawns. In the lab, pyrethroids were consistently several orders of magnitude more toxic than organophosphate insecticides, and more likely to runoff fields at lethal levels according to modeling data. In the field, we found that *Macrobrachium* prawn survival at 31 water contact sites in the lower basin of the Senegal River where schistosomiasis is endemic was predicted by pyrethroid application rates to nearby crop fields after controlling for abiotic and prawn-level factors. Our findings suggest that widely used pyrethroid insecticides can have strong non-target effects on *Macrobrachium* prawns that are biocontrol agents where 400 million people are at risk of human schistosomiasis. Understanding the ecotoxicology of high-risk insecticides may help improve human health in schistosomiasis-endemic regions undergoing agricultural expansion.

## 1. INTRODUCTION

Globally, an estimated 1/3 of crops are produced using pesticides (Zhang et al., 2011). The growing use of synthetic chemicals, including pesticides, is a driver of global change (Bernhardt et al., 2017; MA, 2005) that is outpacing other drivers, such as increasing atmospheric CO_2_ and the loss of both habitat and biodiversity (Bernhardt et al., 2017). The use of agricultural pesticides is expected to increase by 2-5 fold by 2050, particularly in Africa where the human population will double to > 2 billion by 2050 (Tilman et al., 2011; UN, 2020). Pesticides increased harvest yield by approximately one-third across several sub-Saharan African countries (Sheahan et al., 2017). Insecticides are pesticides applied to protect crops from insect damage, with organophosphates and pyrethroids most commonly used by small scale farmers in Africa (Atwood and Paisley-Jones, 2017; Salami et al., 2010) and globally (Zhang, 2018). Organophosphates are a class of insecticides that act by inhibiting the enzyme acetylcholinesterase (Newman and Unger, 2003), whereas pyrethroids interfere with voltage-gated sodium channels (Soderlund and Bloomquist, 1989). Despite different modes of action, widespread use, and benefits for food production, both insecticide classes can be poisonous to humans by contact (Bertrand, 2019), and toxic to aquatic animals (Halstead et al., 2015). Unfortunately, the indirect ecological impacts of pesticides are far less studied than other agents of global change, and adverse effects of pesticides found under laboratory conditions remain largely unverified in the environment (Bernhardt et al., 2017). Additionally, such non-target effects could be influencing aquatic species involved in the transmission of certain infectious diseases of humans (Rohr et al., 2019), including snail-borne human schistosomiasis.

Two genera of freshwater snails in Africa, *Bulinus* and *Biomphalaria*, are responsible for transmitting human schistosomes, parasites that infect more than 250 million people worldwide, nearly 200 million in sub-Saharan Africa, and causes approximately 200,000 deaths in Africa annually (Adenowo et al., 2015; Vos et al., 2016). While many countries are making progress towards elimination of schistosomiasis, disease control in Africa has been hampered by limited access to clean water and sanitation and inconsistent availability of schistosomiasis treatment, among other causes (Grimes et al., 2014). In many African countries, collecting water for drinking and washing household items at local, and sometimes polluted, lakes and rivers is a daily part of life. Thus, individuals receiving drug treatment for the disease can become rapidly re-infected by intermediate host snails that remain in the water and subsequently produce thousands of free-swimming parasites that are released into the water each day for up to a year (Mutuku et al., 2014). Snail abundance, and factors that influence it are key to disease control (Civitello et al., 2022; Civitello et al., 2018; Haggerty et al., 2020; King and Bertsch, 2015; Nguyen et al., 2021; Sokolow et al., 2018; Wood et al., 2019). Importantly, schistosomiasis is especially prevalent in rural areas where agricultural expansion has occurred (Rohr et al., 2019), suggesting that agrochemical pollution of waterways might be one important factor contributing to disease risk. Recent experimental evidence suggests that insecticide runoff into aquatic systems can foster snail populations, and potentially *Schistosoma* transmission risk, by killing important snail predators (Haggerty et al., 2022; Halstead et al., 2018), although the degree to which this happens has yet to be determined.

*Macrobrachium* prawns are natural predators of freshwater snail where half the global human population at risk of schistosomiasis lives (Sokolow et al., 2017). Prawns are also highly sensitive to insecticides (Bajet et al., 2012; Hoover et al., 2020). The river prawns *M. vollenhovenii* and *M. rosenbergii*, native to Africa and the Indo-Pacific region, respectively, are both key invertebrate predators of snails that transmit schistosomiasis (Sokolow et al., 2014) and have been proposed as agent of biological control for schistosomiasis (Ozretich et al. 2022; Sokolow et al. 2015). While *M. rosenbergii* is not native to Africa, it is physiologically very similar to native *M. vollenhovenii* (Savaya-Alkalay et al., 2018) and has already been successfully introduced into parts of Africa within aquaculture facilities (New and Valenti, 2000). Monosex *M. rosenbergii* that are unable to interbreed with *M. vollenhovenii* (Savaya-Alkalay et al., 2018) are biological control agents for schistosomiasis in Africa (Levy et al., 2019). Overall, *M. rosenbergii* has a very similar body size, shape, and claw morphology as *M. vollenhovenii* (Savaya-Alkalay et al., 2018). Thus, if one or both *Macrobrachium* prawns are vulnerable to insecticides, they may represent an important link in the relationship between agricultural expansion and human schistosomiasis (Hoover et al., 2019). Insecticide impacts to *M. vollenhovenii* are important to fully understand declines of native prawns where schistosomiasis is increasing (Sokolow et al., 2017) and because of efforts to introduce the species into African waterways as a public health intervention to control schistosomiasis (Hoover et al., 2019). Recent experimental work has demonstrated that environmentally common insecticide concentrations reduce survival of invertebrate snail predators, including the crayfish *Procambarus alleni* (Halstead et al., 2015) and *Macrobrachium lar* from the Philippines (Bajet et al., 2012). *P. alleni* is a crayfish that is ecologically and morphologically similar to *Macrobrachium*, having two chelae or claws used for foraging on plants or animals, and that was previously used in several mesocosm experiments examining the effects of insecticides on snails and *Schistosoma* parasites (Halstead et al., 2015; Halstead et al., 2018). However, it is unclear whether insecticides used by rural communities in developing countries are reducing the survival of the two *Macrobrachium* species that are among the most important biological control agents of human schistosomiasis.

This study aimed to address the above knowledge gaps by using a laboratory study to determine the relationship between concentrations of six insecticides, three organophosphates and three pyrethroids, and survival of *M. rosenbergii*. We then performed an observational field study using caged *M. vollenhovenii* placed into 31 waterways in Senegal, Africa that varied in organophosphate and pyrethroid applications in their surrounding landscape. Based on previous work using similar invertebrate snail predators and insecticides (Bajet et al., 2012; Halstead et al., 2015), we hypothesized that, in both the laboratory and the field, each insecticide class would lower *Macrobrachium* survival relative to controls, but that pyrethroids would be associated with greater mortality than organophosphates.

## 2. MATERIALS AND METHODS

### 2.1. Lab Study

#### 2.1.1. Selecting test concentrations using an estimated environmental concentration model

Three organophosphate (chlorpyrifos, malathion, and terbufos) and three pyrethroid insecticides (esfenvalerate, λ-cyhalothrin, and permethrin) were selected for this study. Technical grade insecticides were used for all laboratory trials (purity >98%; Chemservice, West Chester, PA, USA). To select experimental concentrations for each chemical we followed the methods of Halstead et al. (2015). Briefly, we used United States Environmental Protection Agency (US EPA) Surface Water Calculator software to generate 150 simulated annual peak estimated environmental concentrations (EECs) in a standardized lentic waterbody that is a set distance from a site where the insecticide was of applied on a standardized crop in accordance with instructions on the product label. The software calculates an EEC based on inputs of pesticide traits (e.g. half-life and adsorption), application amount and frequency, and soil and climatic characteristics (based on a region of interest; see Table S1 for details on the parameters used in the model). In risk assessments conducted by the US EPA, EECs are compared against toxicity values (e.g. LC50) to characterize the likelihood of toxicity at a given level of exposure (USEPA, 2004; USEPA, 1992; Rohr et al. 2016). Evaluation of EECs in this way informs the development of environmental standards, policies, guidelines, and regulations, as well as the registration and reregistration of chemicals for legal use (USEPA, 2004). We took the approach of estimating peak field concentrations because (1) they are used in registration and regulation decisions, (2) environmental concentrations of pesticides in West Africa are rare and thus likely miss peak concentrations (but see Anderson et al., 2014; Diop et al., 2016; Savaya-Alkalay et al., 2018), and (3) studies have shown that field concentrations often do not capture peak concentrations within or across years (Rumschlag et al., 2019a). Importantly, a survey of pesticide residues on vegetables in Senegal revealed pesticide detection frequencies consistent with high frequencies of use, contamination levels that exceeded local regulations, the use of banned pesticides, and use patterns that were inconsistent with label instructions (Diop et al., 2016), suggesting that EECs could be conservative estimates of true environmental concentrations in Senegal. Once a defensible range of EECs was obtained, we selected a range of experimental concentrations (Table S2) for each insecticide that spanned the EECs as well as known LC_50_ values of related species for these or related insecticides (Table S3).

#### 2.1.2. Experimental design

To perform a dose-response study of the six insecticides, we procured juvenile *M. rosenbergii* (25 – 40 mm) from a commercial supplier (Aquaculture of Texas, Inc., Weatherford, TX, USA). Because they were juveniles, they did not yet have secondary sexual characteristics for gender identification. Unfortunately, we could not locate a similar commercial supplier of *M. vollenhovenii* in the United States.

Our LC_50_ experiment used a static, nonrenewal (no water changes) dose-response design with five concentrations for each insecticide (*n* = 5 prawns per concentration, see Table S2 and Fig. 1 for tested concentrations) 10 solvent controls (12 mL/L acetone), and 10 water controls. The concentrations were determined based on a serial dilution of a stock solution (measured concentrations of stock solutions were within 15% of the nominal concentrations based on Abraxis ELISA test kits). Each insecticide tested, including controls, had a total of 45 experimental units (*n* = 45 prawns per insecticide). Prawns were randomly assigned to treatments that were known to the investigator. Our sample size was adopted from a similar experimental study by Halstead et al. (2015), which investigated crayfish but not *Macrobrachium* prawns. We chose a static exposure because preliminary visits to field sites in 2019 using a JDC Instruments Flowatch, water flow meter, detected zero water flow within our water access field sites. We conducted trials in March 2017. Each replicate consisted of a single *M. rosenbergii* in a 500 mL glass jar, filled with 400 mL of artificial spring water (Table S4; (Cohen and Neimark, 1980)), and capped with a screen and maintained in a laboratory at 23.5°C. The water had a pH of 7.7 and starting dissolved oxygen >60%. Each individual was fed 0.16 g of shrimp pellets (Cobalt International, South Carolina, USA) *ad libitum*. Survival was assessed 3, 12, and 24 hours after insecticide application, and daily thereafter for 10 days.

**Figure 1.**
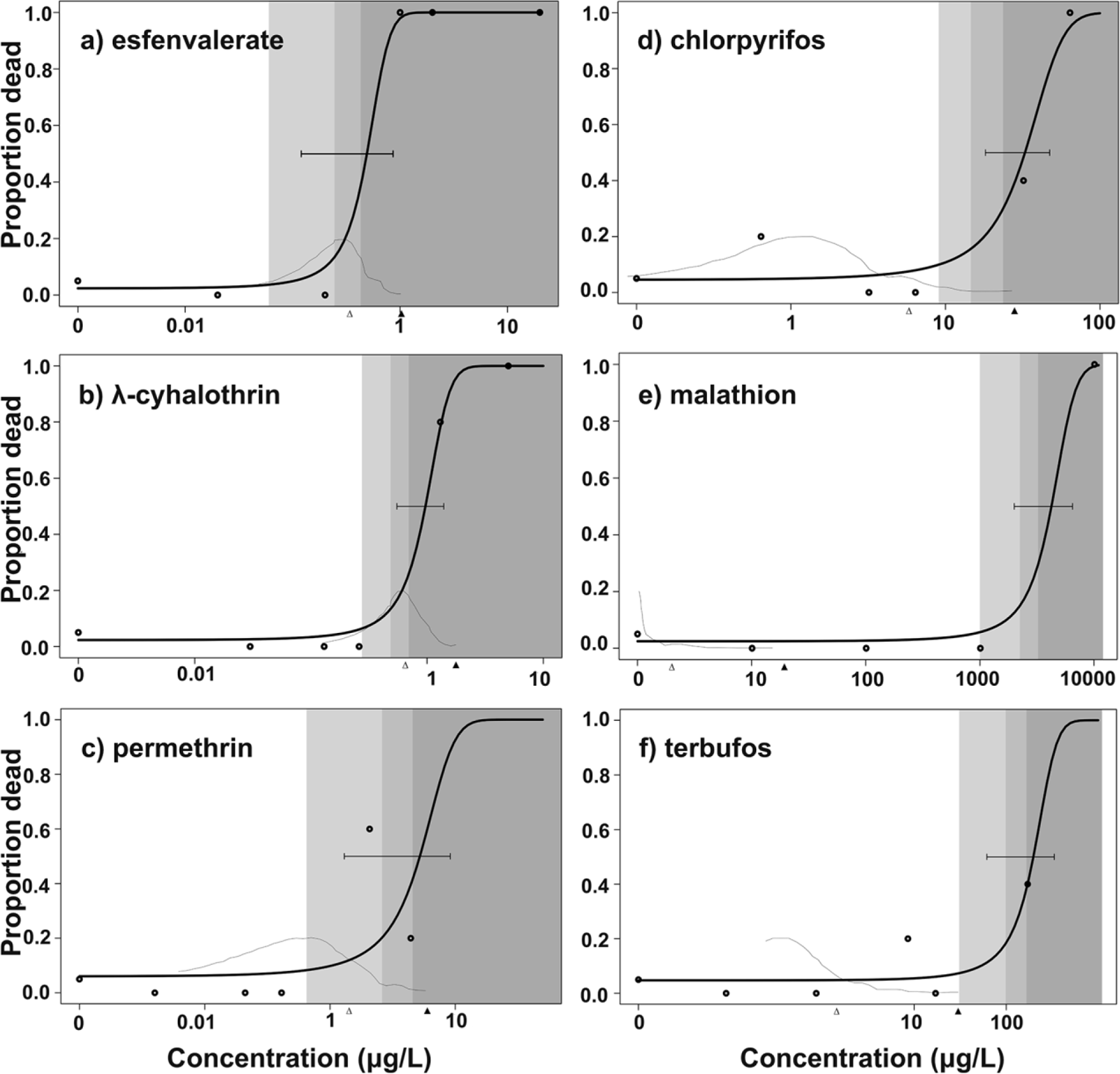
Dose-response curves for *Macrobrachium rosenbergii* after 96-h of exposure to three pyrethroid (a-c) and three organophosphate (d-f) insecticides. The horizontal bar represents the 95% confidence interval around the LC_50_ estimate, with the estimate itself at the point where the confidence interval intersects the curve. The shaded areas represent concentrations above the US EPA’s level of concern of 0.5 x LC_50_ (medium gray) for acute high risk to aquatic organisms. The light gray and dark gray regions represent the area of concern calculated from the lower and upper 95% confidence limits of the LC_50_ estimate, respectively. The gray line curves give the kernel density estimates from 150 simulated annual peak environmental concentrations (EECs) in ponds determined from the US EPA Surface Water Calculator (SWCC) for each insecticide. Thus, those portions of the gray curve within the shaded areas of each plot indicate simulated peak EECs above the US EPA’s level of concern (proportions provided in Table 1). The open and black triangles along the x-axes indicate the median and maximum EECs, respectively, from the SWCC simulations.

#### 2.1.3. Data analyses

We used the *drc* package (Ritz et al., 2015) in *R* 4.0.4 statistical software (RCoreTeam, 2018) to generate dose-response curves and estimate LC_50_ values. Two-parameter logistic models were used to estimate 96-h and 10-d LC_50_ values, and we approximated 95% confidence intervals around each LC_50_ value using the variance of the estimate and then back-transforming from the log scale (Ritz et al., 2015). We then used our LC_50_ estimates to determine the proportion of the simulated EEC values that exceeded the US EPA’s level of concern, which is defined as ≥ 50 percent of the LC_50_ value (0.5 x LC_50_)(USEPA, 2020). Insecticides with EECs that exceeded the level of concern are more likely to enter water bodies at levels toxic to the focal species when applied at their label rates (see Table S1 for label rates).

To determine if differences in prawn survival were more associated with either individual chemicals or chemical class, we performed a Cox mixed effects model for the survival of prawns across all treatments using the R package s*urvival* (Therneau, 2020). We converted all concentrations to toxic units (TUs) using SPEAR Calculator software (v0.8.1, Department System Ecotoxicology – Helmoltz Center for Environmental Research, 2014) to account for variation in absolute toxicity among chemicals, as described by Halstead et al. (2015). Standardized chemical concentration was used as a continuous fixed effect in the model, and random intercepts for each chemical were nested within their respective chemical classes (pyrethroid or organophosphate). Coefficients of the random effects were used to determine the contribution of each chemical and chemical class to overall mortality risk.

### 2.2. Field Study

#### 2.2.1. Study area and village selection

Our field study took place at 31 water points across 16 villages in Northern Senegal, a schistosomiasis hyper-endemic region experiencing rapid agricultural expansion. All of our sites were located along the Senegal and Lampsar Rivers and the shore of Lac de Guiers (16°15′N 15°50′W). Our study region was once populated by *M. vollenhovenii* before the construction of the Diama dam that prevented prawn breeding migrations to estuaries, which led to the loss of prawns upstream of the dam (Savaya Alkalay et al., 2014). Shortly after the dam was constructed, and its associated environmental changes materialized, *Schistosoma* infection increased and led to perennially high infection levels (Steinmann et al., 2006; Talla et al., 1992; Talla et al., 1990).

#### 2.2.2. Insecticide use

We conducted a survey of 663 households at the 16 study villages in 2016 to collect data on the area of crop land where different types of insecticides were used. A respondent from each household was asked to report the area of cultivated land they controlled as well as their use of insecticide on their land. We then calculated the total area on which each class of insecticide was applied in each village. Household surveys were approved by Internal Review Board of the University of California, Santa Barbara (Protocol # 19-170676) and in Senegal by the National Committee of Ethics for Health Research from the Republic of Senegal (Protocol #SEN14/33).

#### 2.2.3. Prawn survival

To investigate prawn survival, we captured wild *M. vollenhovenii* downstream of the Diama dam and temporarily housed them in an outdoor freshwater pond located nearby in St. Louis, Senegal. A subset of mature prawns was collected from the holding pond and transported to experimental cage enclosures at village water points. The average weight (± 1 SD) of prawns used in the experiment was 23.3 g (±9.2). We released adult *M. vollenhovenii* individually into small known-fate enclosures made of a fishing-net overlaid upon a metal frame that was approximately 30 x 30 x 60 cm in size. The cages were not baited and thus prawns were allowed to forage upon prey that naturally entered the cage over time. We deployed one cage for each of 31 water access points across 16 villages. We used a small sample size of 16 villages because of the logistics of performing a village-level social survey and the considerable distance and difficulty of traveling between rural villages in our study region of Senegal. We randomly assigned one prawn to each cage and prawn survival was checked daily by a local villager, given personal protective equipment (waders and shoulder length gloves), who was blind to our agricultural surveys or abiotic conditions near cages. Prawns that died were replaced as needed at the end of each month from the start of the study in March until the end in October 2019 (8 months). Each prawn was one experimental unit (*n* = 225). Before each prawn was placed in a cage, we recorded its mass (g), sex, and number of claws. We also recorded the water temperature (°C) at the cage during the prawn release and various other abiotic and biotic variables (see below).

#### 2.2.4. Estimated runoff and abiotic conditions

We flew a drone with a 12.4 MP camera above each water access point in July 2019 to estimate aquatic vegetation cover because it is very dense in our study region and might intercept insecticide runoff or change abiotic conditions near prawn cages. The drone flew at an altitude of approximately 150 m above the prawn cages, and travelled in all four cardinal directions from those locations, capturing images every 5 seconds to a distance of 300 m from the water access point. All images for each village were aligned using Agisoft Photoscan Professional to create both an orthomosaic and a digital elevation model (DEM). We used the orthomosaic (16 cm/pix resolution) in QGIS 3.2 to estimate the amount of emergent vegetation within a 100-m buffer around each prawn cage. To characterize site topography, we used the DEM (6 m/pix resolution) in QGIS 3.2 to estimate percent slope from each prawn cage to the nearest planted field. Finally, we took the average values of each predictor per village because insecticide use was only available at the village-level. To compare abiotic conditions of sites descriptively in terms of their suitability for prawns, we also visited a random point at each waterway in July 2019 to record average salinity and dissolved oxygen values at each village using a YSI Pro multimeter.

#### 2.2.5. Data analyses

All statistical analyses were conducted with *R* 4.0.4 statistical software (RCoreTeam, 2018). To predict prawn survival, we performed a Cox proportional hazards regression analysis using the *survival* package in *R* and including a single survival time value for each prawn (*n* = 225 total prawns). A Cox model estimates regression coefficients for each predictor and exponentiated coefficients or hazard ratios. A positive regression coefficient suggests that mortality increases with a given predictor. Hazard ratios give the effect sizes, with values > 1, < 1, and equal to 1 representing predictors that increase, decrease, or have no effect on mortality, respectively. Additionally, 95% confidence intervals can be estimated for the hazard ratios. One assumption of the Cox model is that risk is consistent throughout the study. However, that assumption is unlikely to be true in this system because insecticide exposure will temporally vary with rainfall and application times. Unfortunately, precipitation data was not available from our study region to include it as a covariate in our models. Instead, given that exploratory data analyses showed that prawn survival was dependent upon month, we fit a stratified Cox model with a strata term for month, which fits separate baseline hazard functions for each month. Regression coefficients (and associated hazard ratios) optimized for all strata are then fitted. In a stratified Cox model, it is not possible to test for differences among levels of the strata term (here month). The term *+cluster(village)* was included to account for clusters of correlated observations at the village-level (re-sampling of the same villages for prawn survival) and produce robust estimates (standard errors adjusted for the non-independence) using the grouped jackknife method. We fit an initial global model using all village- and prawn-level predictors mentioned above, and a reduced model that sequentially dropped the least significant predictors until all terms were significant. We also tested for an interaction between emergent plant cover and dissolved oxygen, given that dense emergent plants can lower oxygen (Bunch et al., 2010; 2015).

## 3. RESULTS

### 3.1. Lab Study

We found that pyrethroid insecticides were generally an order of magnitude more toxic (lower estimated LC_50_values) than organophosphate insecticides (Table 1). Overall, the LC_50_ (95% CI) of the most toxic pyrethroid and organophosphate were 0.25 μg/L (0.07 - 0.43) for esfenvalerate and 16.73 μg/L (7.86 - 25.60) for chlorpyrifos, respectively (Table 1). Greater toxicity of pyrethroid compared to organophosphate insecticides for *M. rosenbergii* in our laboratory experiment is consistent with data from the US EPA’s Ecotox database for other *Macrobrachium* species (Table 1, Table S3).

**Table 1.**
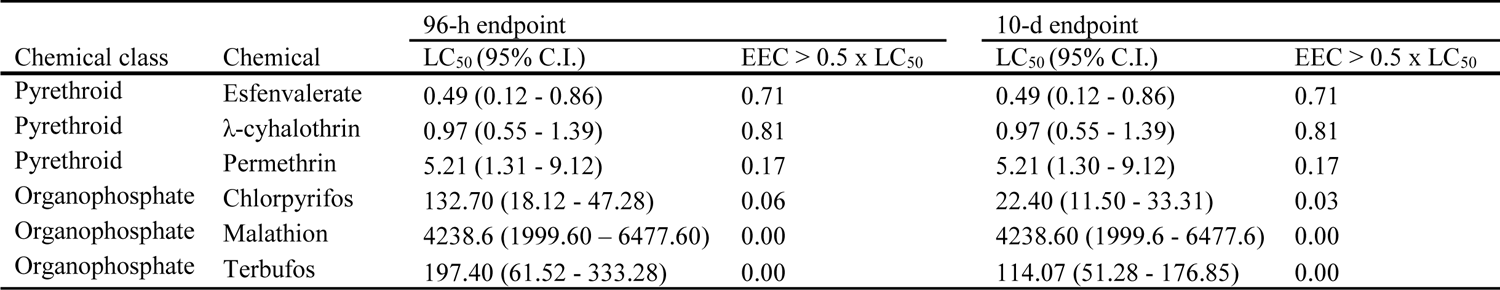
LC_50_ (μg/L) values for *M. rosenbergii* after 96-h and 10-d exposure using all nominal concentrations of three pyrethroid (esfenvalerate, λ-cyhalothrin, and permethrin) and three organophosphate (chlorpyrifos, malathion, and terbufos) insecticides. The second column for each endpoint reports the proportion out of 150 annual peak estimated environmental concentrations (EEC) calculated from the US EPA Pesticide in Water Calculator (v.1.52) that exceeded the US EPA level of concern defined as one-half the estimated LC_50_ (see Methods for full explanation of metrics).

| Chemical class | Chemical | 96-h endpoint |  | 10-d endpoint |  |
| --- | --- | --- | --- | --- | --- |
|  |  | LC<sub>50</sub> (95% C.I.) | EEC > 0.5 x LC<sub>50</sub> | LC<sub>50</sub> (95% C.I.) | EEC > 0.5 x LC<sub>50</sub> |
| Pyrethroid | Esfenvalerate | 0.49 (0.12 - 0.86) | 0.71 | 0.49 (0.12 - 0.86) | 0.71 |
| Pyrethroid | λ-cyhalothrin | 0.97 (0.55 - 1.39) | 0.81 | 0.97 (0.55 - 1.39) | 0.81 |
| Pyrethroid | Permethrin | 5.21 (1.31 - 9.12) | 0.17 | 5.21 (1.30 - 9.12) | 0.17 |
| Organophosphate | Chlorpyrifos | 132.70 (18.12 - 47.28) | 0.06 | 22.40 (11.50 - 33.31) | 0.03 |
| Organophosphate | Malathion | 4238.6 (1999.60 – 6477.60) | 0.00 | 4238.60 (1999.6 - 6477.6) | 0.00 |
| Organophosphate | Terbufos | 197.40 (61.52 - 333.28) | 0.00 | 114.07 (51.28 - 176.85) | 0.00 |

We found that EEC values of each pyrethroid tested commonly exceeded the EPA’s level of concern, defined as half the LC_50_ value (Fig. 1). In contrast, we found that organophosphates rarely exceeded the EPA’s level of concern for *M. rosenbergii* (Table 1). Among organophosphates, only chlorpyrifos generated EEC simulations that exceeded the EPA’s level of concern, which spanned only three percent of the simulations (Table 1). For the three pyrethroids, 17-81% of the exposure simulations exceeded the EPA’s level of concern (Table 1). Thus, pyrethroids had a consistently greater chance of exceeding levels of concern than organophosphates (Fig1; Table 1).

A Cox mixed-effects survival model indicated that insecticide class accounted for 70.7% of the variance in prawn mortality. After 96 hours of exposure, the three most deadly insecticides (i.e., highest hazard ratios or risk per μg/l) were the pyrethroids esfenvalerate and λ-cyhalothrin, and the organophosphate chlorpyrifos (Table S5, Table 2). Converting all concentrations to toxic units (TUs) using the SPEAR Calculator software revealed that, on average, pyrethroid insecticides led to 275% more mortality than organophosphates (Table S5). The coefficients of random effects suggest that variation among individual insecticides within the organophosphate class was largely driven by the relatively low risk presented by malathion (Table 1 and 2).

**Table 2.** Cox survival analysis for *Macrobrachium rosenbergii* exposed to multiple concentrations of three pyrethroid (esfenvalerate, λ-cyhalothrin, and permethrin) and three organophosphate (chlorpyrifos, malathion, and terbufos) insecticides for 10 days. Positive coefficients (coef) indicate that the probability of prawn mortality during the study increased with chemical exposure. The hazard ratio (HR) is the exponent of the coefficient and indicates the probability of an increase in mortality for every 1 µg/L increase in concentration. For example, the hazard ratio of 1.051 for esfenvalerate indicates that every 1 µg/L increase in esfenvalerate increases the probability of mortality during the study increases by 5.1%. The 95% confidence intervals are provided for the hazard ratio.

| Chemical | *coef* | *SE* | *HR (95%CI)* | *z-value* | *p-value* |
| --- | --- | --- | --- | --- | --- |
| Esfenvalerate | 0.05 | 0.01 | 1.05 (1.04-1.07) | 6.62 | <0.001 |
| λ-cyhalothrin | 0.63 | 0.09 | 1.87 (1.69-2.06) | 6.69 | <0.001 |
| Permethrin | 0.29 | 0.16 | 1.34 (1.02-1.67) | 1.80 | 0.072 |
| Chlorpyrifos | 0.16 | 0.02 | 1.17 (1.12-1.23) | 7.02 | <0.001 |
| Malathion | 0.00 | 0.00 | 1.00 (1.00-1.00) | 6.42 | <0.001 |
| Terbufos | 0.01 | 0.00 | 1.02 (1.01-1.02) | 4.60 | <0.001 |

### 3.2. Field Study

We documented a total of 1,515 ha of agricultural fields using our social survey in the 16 villages we sampled in Senegal (Table 3). Insecticides were applied to 60% of the total planted field area and there was an average of 47.8 ha (± 8.0 SE) of insecticide application per village. One organophosphate (dimethoate) and one pyrethroid (deltamethrin) insecticide together made up 78% of the total area where insecticides were applied. Each village received an average of 14.1 prawns (± 1.4 SE) during the study, which depended upon number of water access points at each village.

**Table 3.** Results of the Cox proportional hazards model after model selection (see methods for selection process). Average values for all predictors in the initial model are provided in Table S6.

| Predictor | *coef* | *robust SE<sup>a</sup>* | *hazard (95% CI)* | *z-value* | *p-value* |
| --- | --- | --- | --- | --- | --- |
| Total pyrethroid use (ha) | 0.06 | 0.01 | 1.06 (1.03-1.09) | 4.11 | <0.001 |
| Water temperature (°C) | 0.31 | 0.10 | 1.36 (1.12-1.64) | 3.16 | 0.002 |
| Sex of prawn (male) | 0.43 | 0.19 | 1.53 (1.05-2.24) | 2.20 | 0.028 |
| Avg. emergent vegetation (m<sup>2</sup>) | <0.01 | <0.01 | 1.00 (1.00-1.00) | 3.92 | <0.001 |
| Avg. dissolved oxygen (ppm) | -0.19 | 0.05 | 0.83 (0.76-0.90) | -4.23 | <0.001 |
| Avg. conductivity (µS/cm) | <0.01 | <0.01 | 1.01 (1.00-1.02) | 3.13 | 0.002 |

<sup>a</sup>Wald tests were substituted for likelihood ratio tests to provide robust variances.

Given that *Macrobrachium* spp. LC_50_ values for pyrethroids were one to two orders of magnitude lower than LC_50_ values for organophosphates and the laboratory hazard ratios suggested that pyrethroids were generally more toxic in nature than organophosphates, we hypothesized that pyrethroid use (measured as area of insecticide application) would be more positively associated with prawn mortality in Senegalese waterbodies than organophosphate use. As predicted, when accounting for significant covariates in the final model (Table 3), *M. vollenhovenii* mortality was positively associated with total pyrethroid applications (ha) (Fig. 2a), but was not significantly related to organophosphate applications (Table S5, Table 3). In villages with pyrethroid use, prawn survival decreased rapidly in the first few days after prawn release, consistent with the 96-h results of the LC_50_ trials.

**Figure 2.**
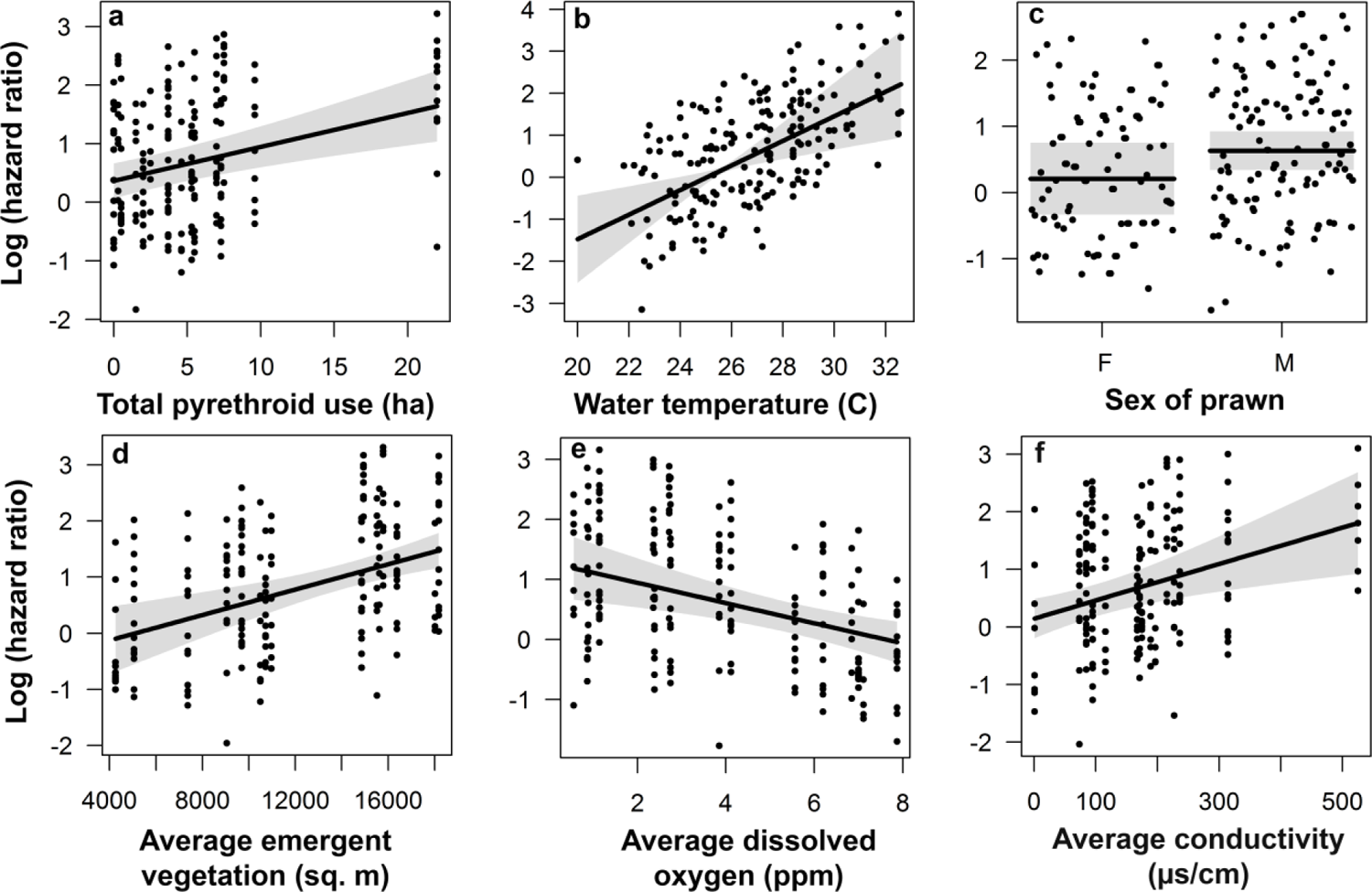
Partial residual plot of the final Cox model (after model selection) showing the association between prawn mortality (log(hazard ratio)) and pyrethroid use (a), water temperature (b), sex of the prawn (c), average emergent vegetation with 100m of the prawn cage (d), average dissolved oxygen (e), and average conductivity (f). Each panel shows the marginal effect controlling for the other factors in the model, which are shown in the remaining panels. Plots were generated in the R package *visreg*. Shaded areas represent 95% confidence intervals.

Prawn mortality was also associated with several covariates. For example, prawn mortality was positively associated with water point temperature at release (Table 3; Fig. 2b; mean water temperature: 28.2° C, range: 20 - 32.6° C), and the average amount of emergent vegetation within 100 m of cages (Fig. 2d). Male prawns experienced significantly higher mortality than female prawns (Table 3; Fig. 2c). Finally, prawn mortality was negatively associated with dissolved oxygen (Fig. 2e), whereas mortality was positively associated with conductivity (Fig. 2f). We found no significant interaction between emergent plant cover and dissolved oxygen (Table S5; *p* > 0.05).

## 4. DISCUSSION

Synthetic chemicals are important agents of global change, but adverse effects of pesticides observed in laboratory studies are rarely estimated in the field (Bernhardt et al., 2017). Our laboratory and field experiments support the hypothesis that pyrethroid insecticides pose a high mortality risk for two species of *Macrobrachium* predators of snails whose declines have been positively linked to schistosomiasis infections in humans (Sokolow et al., 2017). We found that *M. rosenbergii* LC_50_ values for three pyrethroids were consistently lower by one or more orders of magnitude than organophosphate insecticides, in agreement with previous laboratory studies of other invertebrate snail predators (Bajet et al., 2012; Halstead et al., 2015). Importantly, expected concentrations of pyrethroid insecticides in water bodies are more likely to exceed LC_50_ values of *Macrobrachium* prawns than expected field concentrations of organophosphate insecticides. Thus, our findings suggest that in nature, pyrethroid insecticides may be more likely to runoff fields and kill prawns than organophosphate insecticides. We corroborated these laboratory findings in a natural setting by documenting that caged *M. vollenhovenii* survival at water points was best predicted by reported pyrethroid rather than organophosphate applications on crop fields near these water points. Thus, prawn survival was negatively associated with pyrethroid and not organophosphate use, despite our study sites actually averaging more total organophosphate than pyrethroid insecticide applications reported by households living nearby. Although the pyrethroid and organophosphates insecticides used in the field did not perfectly match those studied in the laboratory, in several studies, toxicity levels were more similar within than across pesticides classes (Rumschlag et al., 2022; Rumschlag et al., 2019b; Rumschlag et al., 2021). Consequently, our findings suggest that predicted rises in insecticide use associated with human population growth (Tilman et al., 2011) could be highly relevant to interventions employing *Macrobrachium* prawns.

Introducing *Macrobrachium* prawns into waterways has recently been proposed as a public health intervention in our study region (Sokolow et al., 2017), and identifying abiotic factors that affect prawn survival in the wild will be very important to the success of such interventions. Consistent with previous studies (Cheng et al., 2003), we found that oxygen was a strong positive determinant of *Macrobrachium* survival. Prawn mortality increases quickly below 2 mg/L oxygen (Ferreira et al., 2011), and approximately 25% of our field measurements were below this threshold. Temperature increases the metabolic rate of ectotherms, with optimal temperatures being approximately 30°C for both *M. rosenbergii* and *M. vollenhovenii* (Akinwunmi et al., 2014; New, 1995). Prawns in our experiment were likely acclimated to ambient temperatures in our outdoor growing ponds but could not migrate to more favorable temperatures in waterways once placed in experimental cages. Summer water temperatures in the Senegal River Basin can reach 32°C (Sane et al., 2017), with air temperatures reaching 40°C (Cheikh et al., 2013). Given that each degree Celsius rise in temperatures increases oxygen demand by increasing prawn metabolism (Manush et al., 2004; Xi-lin et al., 1999), water temperatures at release sites may have increased prawn mortality by raising oxygen demands.

Prawn- and village-level characteristics may also influence prawn responses to abiotic conditions in Senegalese waterways. Male *M. vollenhovenii* could be more sensitive to oxygen because they reach a larger size (Olele and Kalayolo, 2012), and body size in crayfish determines oxygen consumption (Armitage and Wall, 1982). Unlike temperature, conductivity does not impact oxygen stress in *M. rosenbergii* (Ern et al., 2013). All conductivity values that we observed were suitable for adult prawns (New, 1995). However, water conductivity is an indicator of agricultural runoff (Harwell et al., 2008). Eutrophication associated with nutrients in runoff can increase the chances of hypoxia at sites (Dodds and Whiles, 2019) and eutrophication from nutrients has been observed in our study system (Cogels et al., 2001). Fertilizers, which are used at all 16 villages, are also positively associated with invasive macrophytes, such as *Typha* spp., that were the most common emergent aquatic plant in our study and are distributed worldwide (Bansal et al., 2019). Emergent plants such as *Typha* spp. can shade waterways, limiting photosynthesis by algae or submerged plants that produce oxygen and change water circulation (Jones et al. 2021). *Typha* spp. can also create leaf litter that lowers dissolved oxygen as it decays (Bunch et al., 2010; 2015). However, we found no significant interaction between emergent plant cover and dissolved oxygen. Although no previous study has, to our knowledge, examined prawns in relation to emergent plants, *Typha* spp. invasion can lead to aquatic communities dominated by hypoxic-tolerant species (Schrank and Lishawa, 2019). Crayfish, which are phylogenetically and functionally similar to prawns, will feed on a variety of aquatic plants but do not readily consume *Typha* (Bolser et al., 1998). This might suggest that *Typha* may also provide little direct benefit to *Macrobrachium*. Together, these findings suggest that prawn- and site-level factors can influence prawn mortality that, in turn, can have important impacts on population densities of intermediate host snails of human schistosomiasis.

Extrapolating hazards among insecticide classes from laboratory to field settings is key to understanding the effects that different insecticide classes might have on *Macrobrachium* biocontrol of schistosomiasis. Our laboratory *M. rosenbergii* LC_50_ values for three organophosphates were generally within the 95% CIs of the LC_50_’s reported for *P. alleni* from Halstead et al. (Halstead et al., 2015). Similar to Halstead et al. (2015), we found that the two insecticides with the lowest hazard ratios were the organophosphates malathion and terbufos, whereas chlorpyrifos posed a higher risk among the organophosphates. Additionally, previous laboratory studies support our finding of greater toxicity of pyrethroid than organophosphate insecticides to crayfish and *Macrobrachium* prawns (Bajet et al., 2012; Halstead et al., 2015; Halstead et al., 2018). Pyrethroids, including deltamethrin, the most common pyrethroid reported in our household surveys in Senegal, had such a high toxicity to *M. lar* in the Philippines that pyrethroid environmental concentrations actually exceeded LC_50_ values in the laboratory (Bajet et al., 2012). Environmental concentrations simulated in our study, using EPA software showed patterns consistent within insecticide class and with previous studies (Halstead et al., 2015). However, we are the first to provide evidence from nature supporting all of these laboratory findings, thus providing evidence that toxicity at relevant exposures (Rohr et al. 2011).

The loss of river prawns in the Senegal River Delta following agricultural projects that coincided with disease outbreaks (Sokolow et al., 2017; Steinmann et al., 2006) emphasizes the need to identify low-risk insecticides for increasing crop yields without harming native prawns. Successfully re-introducing *Macrobrachium* prawns for biocontrol of schistosomiasis will require identifying low-risk insecticides in endemic and developing regions undergoing agricultural expansion. Among organophosphates, we found that malathion has a particularly low toxicity to prawns, consistent with experiments using the prawn species *M. lar* (Bajet et al., 2012) and the crayfish *P. alleni* (Halstead et al., 2015). As the *M. rosenbergii* used in our laboratory study were commercially bred for human consumption in a hatchery, they had no known previous exposure to insecticides in their familial history, which strongly suggests that the displayed resistance by *Macrobrachium* prawns to malathion is innate. Thus, our results suggest that malathion may be a particularly useful insecticide to protect crops from pests without increasing *Macrobrachium* mortality, a conclusion recently supported by freshwater mesocosm studies on schistosomiasis risk more generally (Haggerty et al., 2022). Although the pyrethroids we tested were generally more toxic to prawns than the organophosphates, the pyrethroid permethrin had a lower chance of reaching EPA levels of concern (EEC > 0.5 x LC_50_) than λ-cyhalothrin or esfenvalerate pyrethroids. Permethrin also has the lowest desorption rate among pyrethroids we examined (Fojut and Young, 2011), which could be important for lowering its bioavailability in agricultural regions where organic carbon levels are low (Fojut and Young, 2011).

Our study has several limitations that could influence our understanding of insecticide effects on prawn biocontrol agents in field settings. We did not have spatial information on the agricultural area reported in our social surveys. Thus, we assumed that villages with more fields also have more fields near their water access points. However, some fields reported in our surveys were likely too distant to generate runoff into waterways. In this case, our village-level analyses might not capture the agricultural runoff or abiotic conditions that occur at prawn cages as accurately as if we had temporal abiotic samples or knew each field location in relation to the cages. An additional caveat to our field study is that we did not have insecticide application rate data, and, thus, we assumed that chemical application rates were comparable among the study villages and did not change between our survey and prawn trials. Quantifying chemical concentrations in waterways each month could have helped to address several of the above limitations and to determine if static exposures in the laboratory overestimated *Macrobrachium* exposure to insecticides in nature, but quantifying monthly insecticide concentrations from waterways receiving a potential mixture of chemicals can be logistically challenging and costly. Non-lethal effects of organophosphates that were not measured in our study, appear to occur for *Macrobrachium rosenbergii* (Chang et al., 2013) and may have yet unknown impacts on their long-term use as biocontrol agents near agricultural areas. Future studies that can address the limitations of our field study and that explore known interactions between organophosphates and pyrethroids that can increase toxicity (Gaughan et al., 1980) would be useful to further improve our understanding of insecticide risks.

In conclusion, our findings suggest that levels of different insecticide classes used by rural subsistence farmers near waterways may adversely affect *Macrobrachium* prawn species that are biocontrol agents of schistosomiasis. Importantly, previous mesocosm and modeling studies demonstrated that snails that can transmit trematode parasites are less sensitive to insecticides than arthropod predators and competitors of snails (Becker et al., 2020; Halstead et al., 2015; Halstead et al., 2018; Hoover et al., 2020; Rumschlag et al., 2021) and that the loss of snail predators arising from insecticide toxicity can increase snail densities and possibly the risk of snail-transmitted disease (Halstead et al., 2018; Rohr et al., 2008). Thus, our findings may offer one potential explanation for the positive links between *Schistosoma* transmission and agricultural expansion and can help inform future *Macrobrachium* prawn introductions to control snails. Our findings further suggest that natural recolonization of prawns in aquatic systems may be hampered by insecticide runoff. While insecticides will remain essential in developing countries (Snyder et al., 2015), educating farmers about the risks of particular insecticides (particularly pyrethroids) for native fauna may be warranted. Future studies are needed to examine the effects of farmers’ switching from pyrethroids to alternative insecticides with fewer impacts on arthropod predators of snails, such as malathion (Lund et al. 2021). Careful choice of insecticides may be needed to reduce crop pests without increasing the risk of disease in areas endemic for schistosomiasis.

## Database deposition

Data and code used in this manuscript are available as a Figshare data repository: https://www.doi.org/10.6084/m9.figshare.14331272.v1.

## Author contributions

CJEH, BKD, NJ, and JRR conceived and designed the experiment. PDN, NJ, and GR lead prawn cultivation in Senegal and organized the field project. AJL and DLC conducted village surveys of insecticide use. CJEH and BKD conducted the statistics and generated the figures. CJEH, BKD, JRR wrote the initial manuscript draft of the field and lab experiment and all authors contributed to the preparation of the manuscript.

## Acknowledgements

The authors declare no competing financial interest. We are grateful for the staff of Espoir Pour La Santé for their cooperation in the field aspects of the project, villagers who monitored the caged prawns, and to Amit Savaia of Ben-Gurion University for assistance with prawn transport and cage methodological advice.

## References

1. Adenowo, A.F., Oyinloye, B.E., Ogunyinka, B.I., et al. 2015. Impact of human schistosomiasis in sub-Saharan Africa. Braz J Infect Dis 19(2), 196–205.

2. Akinwunmi, M.F., Bello Olusoji, O.A. and Sodamola, M.Y. 2014. The rearing of African river prawn, Macrobrachium vollenhovenii in concrete tank using locally formulated diet. International Journal of Fisheries and Aquatic Studies 2(2), 265–270.

3. Anderson, K.A., Seck, D., Hobbie, K.A., et al. 2014. Passive sampling devices enable capacity building and characterization of bioavailable pesticide along the Niger, Senegal and Bani Rivers of Africa. Philosophical Transactions of the Royal Society B: Biological Sciences 369(1639), 20130110.

4. Armitage, K.B. and Wall, T.J. 1982. The effects of body size, starvation and temperature acclimation on oxygen consumption of the crayfish Orconectes nais. Comparative Biochemistry and Physiology Part A: Physiology 73(1), 63–68.

5. Atwood, D. and Paisley-Jones, C. 2017 Pesticides industry sales and usage: 2008–2012 Market Estimates, US Environmental Protection Agency, Washington, DC.

6. Bajet, C.M., Kumar, A., Calingacion, M.N., et al. 2012. Toxicological assessment of pesticides used in the Pagsanjan-Lumban catchment to selected non-target aquatic organisms in Laguna Lake, Philippines. Agricultural Water Management 106, 42–49.

7. Bansal, S., Lishawa, S.C., Newman, S., et al. 2019. Typha (Cattail) Invasion in North American Wetlands: Biology, Regional Problems, Impacts, Ecosystem Services, and Management. Wetlands 39(4), 645–684.

8. Becker, J.M., Ganatra, A.A., Kandie, F., et al. 2020. Pesticide pollution in freshwater paves the way for schistosomiasis transmission. Scientific reports 10(1), 1–13.

9. Bernhardt, E.S., Rosi, E.J. and Gessner, M.O. 2017. Synthetic chemicals as agents of global change. Front. Ecol. Environ. 15(2), 84–90.

10. Bertrand, P.G. (2019) Pesticides use and misuse and their impact in the environment. Larramendy, M.L. (ed), IntechOpen, London, UK.

11. Bolser, R.C., Hay, M.E., Lindquist, N., et al. 1998. Chemical defenses of freshwater macropytes against crayfish herbivory. Chemical Ecology 24(10), 1639–1658.

12. Bunch, A.J., Allen, M.S. and Gwinn, D. 2010. Spatial and temporal hypoxia dynamics in dense emergent macrophytes in a Florida lake. Wetlands 30(3), 429–435.

13. Bunch, A.J., Allen, M.S. and Gwinn, D. 2015. Influence of macrophyte-induced hypoxia on fish communities in lakes with altered hydrology. Lake and reservoir managment 31, 11–19.

14. Chang, C.C., Rahmawaty, A. and Chang, Z.W. 2013. Molecular and immunological responses of the giant freshwater prawn, Macrobrachium rosenbergii, to the organophosphorus insecticide, trichlorfon. Aquat Toxicol 130–131, 18-26.

15. Cheikh, B.G., Moctar, D. and Raymond, M. 2013. Assessing the impacts of climate change on water resources of a West African trans-boundary river basin and its environmental consequences (Senegal River Basin). Sciences in Cold and Arid Regions 5(1), 140.

16. Cheng, W., Liu, C.-H. and Kuo, C.-M. 2003. Effects of dissolved oxygen on hemolymph parameters of freshwater giant prawn, Macrobrachium rosenbergii (de Man). Aquaculture 220(1), 843–856.

17. Civitello, D.J., Angelo, T., Nguyen, K.H., et al. 2022. Transmission potential of human schistosomes can be driven by resource competition among snail intermediate hosts. P. Natl. Acad. Sci. USA 119(6), 9.

18. Civitello, D.J., Fatima, H., Johnson, L.R., et al. 2018. Bioenergetic theory predicts infection dynamics of human schistosomes in intermediate host snails across ecological gradients. Ecol. Lett. 21(5), 692–701.

19. Cogels, F.X., Frabouiet-Jussiia, S. and Varis, O. 2001. Multipurpose use and water quality challenges in Lac de Guiers (Senegal). Water science and technology: a journal of the International Association on Water Pollution Research 44(6), 35–46.

20. Cohen, L. and Neimark, H. 1980. Schistosoma mansoni:response of cercariae to a thermal gradient. Parasitology 66, 362–364.

21. Diop, A., Diop, Y.M., Thiare, D.D., et al. 2016. Monitoring survey of the use patterns and pesticide residues on vegetables in the Niayes zone, Senegal. Chemosphere 144, 1715–1721.

22. Dodds, W. and Whiles, M. (2019) Freshwater Ecology, Academic Press, Cambridge, MA.

23. Ern, R., Huong, D.T.T., Nguyen, V.C., et al. 2013. Effects of salinity on standard metabolic rate and critical oxygen tension in the giant freshwater prawn (Macrobrachium rosenbergii). Aquaculture Research 44(8), 1259–1265.

24. Ferreira, N.C., Bonetti, C. and Seiffert, W.Q. 2011. Hydrological and Water Quality Indices as management tools in marine shrimp culture. Aquaculture 318(3), 425–433.

25. Fojut, T.L. and Young, T.M. 2011. Desorption of pyrethroids from suspended solids. Environ Toxicol Chem 30(8), 1760–1766.

26. Gaughan, L.C., Engel, J.L. and Casida, J.E. 1980. Pesticide interactions: effects of organophosphorus pesticides on the metabolism, toxicity, and persistence of selected pyrethroid insecticides. Pestic. Biochem. Phys. 14(1), 81–85.

27. Grimes, J.E., Croll, D., Harrison, W.E., et al. 2014. The relationship between water, sanitation and schistosomiasis: a systematic review and meta-analysis. PLoS Negl Trop Dis 8(12), e3296.

28. Haggerty, C.J.E., Bakhoum, S., Civitello, D.J., et al. 2020. Aquatic macrophytes and macroinvertebrate predators affect densities of snail hosts and local production of schistosome cercariae that cause human schistosomiasis. Plos Neglect. Trop. Dis. 14(7), 25.

29. Haggerty, C.J.E., Halstead, N.T., Civitello, D.J., et al. 2022. Reducing disease and producing food: Effects of 13 agrochemicals on snail biomass and human schistosomes. J. Appl. Ecol. 59(3), 729–741.

30. Halstead, N.T., Civitello, D.J. and Rohr, J.R. 2015. Comparative toxicities of organophosphate and pyrethroid insecticides to aquatic macroarthropods. Chemosphere 135, 265–271.

31. Halstead, N.T., Hoover, C.M., Arakala, A., et al. 2018. Agrochemicals increase risk of human schistosomiasis by supporting higher densities of intermediate hosts. Nat Commun 9(1), 837.

32. Harwell, M.C., Surratt, D.D., Barone, D.M., et al. 2008. Conductivity as a tracer of agricultural and urban runoff to delineate water quality impacts in the northern Everglades. Environmental monitoring and assessment 147(1-3), 445–462.

33. Hoover, C.M., Rumschlag, S.L., Strgar, L., et al. 2020. Effects of agrochemical pollution on schistosomiasis transmission: a systematic review and modelling analysis. Lancet Planet. Health 4(7), E280–E291.

34. Hoover, C.M., Sokolow, S.H., Kemp, J., et al. 2019. Modelled effects of prawn aquaculture on poverty alleviation and schistosomiasis control. Nature Sustainability 2(7), 611–620.

35. Jones I.J., Sokolow S.H., Chamberlin A., Lund A.J., Jouanard N., Bandagny L., Ndione R., Senghor S., Schacht A.M., Riveau G., Hopkins S.R., Rohr J.R., Remais J., Lafferty K.D., Kuris A.M., Wood C.L., De Leo, G.A.. 2021. Schistosome infection in Senegal is associated with different spatial extents of risk and ecological drivers for S. haematobium and S. mansoni. PLoS Neglected Tropical Diseases, 15(9):e0009712, https://doi.org/10.1371/journal.pntd.0009712

36. King, C.H. and Bertsch, D. 2015. Historical perspective: snail control to prevent schistosomiasis. PLoS Negl Trop Dis 9(4), e0003657.

37. Levy, T., Rosen, O., Manor, R., et al. 2019. Production of WW males lacking the masculine Z chromosome and mining the Macrobrachium rosenbergii genome for sex-chromosomes. Scientific Reports 9(1), 12408.

38. Lund A.J., Lopez-Carr, D., Sokolow, S.H., Rohr, J.R., De Leo, G.A. 2021. Agricultural innovations to reduce the health impacts of dams. Sustainability, 13(4), 1869. https://doi.org/10.3390/su13041869.

39. MA, Millenium Ecosystem Assessment 2005 Ecosystems and human well-being: synthesis. Millenium Ecosystem Assessment (ed), Island Press, Washington DC.

40. Manush, S.M., Pal, A.K., Chatterjee, N., et al. 2004. Thermal tolerance and oxygen consumption of Macrobrachium rosenbergii acclimated to three temperatures. Journal of Thermal Biology 29(1), 15–19.

41. Mutuku, M.W., Dweni, C.K., Mwangi, M., et al. 2014. Field-derived Schistosoma mansoni and Biomphalaria pfeifferi in Kenya: a compatible association characterized by lack of strong local adaptation, and presence of some snails able to persistently produce cercariae for over a year. Parasit Vectors 7, 533.

42. New, M.B. 1995. Status of freshwater prawn farming. Aquacul Res 26, 1–54.

43. New, M.B. and Valenti, W.C. (2000) Freshwater prawn culture: the farming of Macrobrachium rosenbergii, pp. 1–10, Blackwell Science, London, England.

44. Newman, M.C. and Unger, M.A. (2003) Fundamentals of Ecotoxicology, CRC Press, Boca Raton, FL.

45. Nguyen, K.H., Boersch-Supan, P.H., Hartman, R.B., et al. 2021. Interventions can shift the thermal optimum for parasitic disease transmission. P. Natl. Acad. Sci. USA 118(11), 8.

46. Olele, N.F. and Kalayolo, P.E. 2012. Morphometric characteristics of the giant African river prawn, Macrobrachium vollenhovenii (Herklot, 1857) caught from Warri River coast. Journal of Agriculture and Biological Sciences 3(1), 232-239.

47. Ozretich, RW., Wood, C.L., Allan, F., Koumi, A.R., Norman, R., Brierley, A.S., De Leo, G.A., Little, D.C., 2022. The potential for aquaculture toreduce poverty and control schistosomiasis in Côte d’Ivoire (Ivory Coast) during an era of climate change: A systematic review. Rev. Fish. Sci. Aquac. 1–31. https://doi.org/10.1080/23308249.2022.2039096

48. R Core Team 2018 R: A language and environment for statistical computing, Vienna, Austria.

49. Ritz, C., Baty, F., Streibig, J.C., et al. 2015. Dose-Response Analysis Using R. PLOS ONE 10(12).

50. Rohr, J.R., Barrett, C.B., Civitello, D.J., et al. 2019. Emerging human infectious diseases and the links to global food production. Nat Sustain 2(6), 445–456.

51. Rohr, J.R., Salice, C.J., Nisbet, R.M. 2016. The pros and cons of ecological risk assessment based on data from different levels of biological organization. Critical Reviews in Toxicology 46(9), 756–784.

52. Rohr, J.R., Sesterhenn, T.M., Stieha, C. 2011. Will climate change reduce the effects of a pesticide on amphibians?: partitioning effects on exposure and susceptibility to contaminants. Global Change Biology 17(2), 657–666.

53. Rohr, J.R., Schotthoefer, A.M., Raffel, T.R., et al. 2008. Agrochemicals increase trematode infections in a declining amphibian species. Nature 455(7217), 1235–1239.

54. Rumschlag, S.L., Bessler, S.M. and Rohr, J.R. 2019a. Evaluating improvements to exposure estimates from fate and transport models by incorporating environmental sampling effort and contaminant use. Water Res.

55. Rumschlag, S.L., Casamatta, D.A., Mahon, M.B., et al. 2022. Pesticides alter ecosystem respiration via phytoplankton abundance and community structure: Effects on the carbon cycle? Glob. Change Biol. 28(3), 1091–1102.

56. Rumschlag, S.L., Halstead, N.T., Hoverman, J.T., et al. 2019b. Effects of pesticides on exposure and susceptibility to parasites can be generalised to pesticide class and type in aquatic communities. Ecol. Lett.

57. Rumschlag, S.L., Mahon, M.B., Hoverman, J.T., et al. 2021. Consistent effects of pesticides on community structure and ecosystem function in freshwater systems (vol 11, 6333, 2020). Nature Communications 12(1), 4.

58. Salami, A., Kamara, A.B. and Brixiova, Z. 2010 Smallholder agriculture in East Africa:Trends, Constraints and Opportunities, African Development Bank, Tunis, Tunisia.

59. Sane, S., Ngansoumana, B., Arfi, R., et al. 2017. Environmental conditions and primary production in a Sahelian shallow lake (Lake Guiers, North Senegal). International Journal of Biological and Chemical Sciences 11(3).

60. Savaya-Alkalay, A., Ndao, P.D., Jouanard, N., et al. 2018. Exploitation of reproductive barriers between Macrobrachium species for responsible aquaculture and biocontrol of schistosomiasis in West Africa. Aquaculture Environment Interactions 10, 487–499.

61. Savaya Alkalay, A., Rosen, O., Sokolow, S.H., et al. 2014. The prawn Macrobrachium vollenhovenii in the Senegal River basin: towards sustainable restocking of all-male populations for biological control of schistosomiasis. PLoS Negl Trop Dis 8(8), e3060.

62. Schrank, A.J. and Lishawa, S.C. 2019. Invasive cattail reduces fish diversity and abundance in the emergent marsh of a Great Lakes coastal wetland. Journal of Great Lakes Research 45(6), 1251–1259.

63. Sheahan, M., Barrett, C. and Goldvale, C. 2017. Human health and pesticide use in Sub-Saharan Africa. Agricultural Economics 48(S1), 27–41.

64. Snyder, J., Smart, J., Goeb, J., et al. 2015 Pesticide use in Sub-Saharan Africa: Estimates, Projections, and Implications in the Context of Food System Transformation.

65. Soderlund, D.M. and Bloomquist, J.R. 1989. Neurotoxic actions of pyrethroid insecticides. Annu Rev Entomol 34, 77–96.

66. Sokolow S.H., Huttinger E., Jouanard J., et al. 2015 Reduced transmission of human schistosomiasis after restoration of a native river prawn that preys on the snail intermediate host. Proceeding of the National Academy of Science. 112(31): 9650–9655.

67. Sokolow, S.H., Jones, I.J., Jocque, M., et al. 2017. Nearly 400 million people are at higher risk of schistosomiasis because dams block the migration of snail-eating river prawns. Philos Trans R Soc Lond B Biol Sci 372(1722).

68. Sokolow, S.H., Lafferty, K.D. and Kuris, A.M. 2014. Regulation of laboratory populations of snails (Biomphalaria and Bulinus spp.) by river prawns, Macrobrachium spp. (Decapoda, Palaemonidae): implications for control of schistosomiasis. Acta tropica 132, 64–74.

69. Sokolow, S.H., Wood, C.L., Jones, I.J., et al. 2018. To Reduce the Global Burden of Human Schistosomiasis, Use ‘Old Fashioned’ Snail Control. Trends Parasitol. 34(1), 23–40.

70. Steinmann, P., Keiser, J., Bos, R., et al. 2006. Schistosomiasis and water resources development: systematic review, meta-analysis, and estimates of people at risk. Lancet Infect Dis 6(7), 411–425.

71. Talla, I., Kongs, A. and Verlé, P. 1992. Preliminary study of the prevalence of human schistosomiasis in Richard-Toll (the Senegal river basin). Transactions of The Royal Society of Tropical Medicine and Hygiene 86(2), 182–182.

72. Talla, I., Kongs, A., Verle, P., et al. 1990. Outbreak of intestinal schistosomiasis in the Senegal River Basin. Annales de la Société belge de médecine tropicale 70, 173–180.

73. Therneau, T. 2020 A package for survival analysis in R.

74. Tilman, D., Balzer, C., Hill, J., et al. 2011. Global food demand and the sustainable intensification of agriculture. Proceedings of the National Academy of Sciences 108(50), 20260.

75. UN 2020 Population, United Nations.

76. USEPA (2004) An Examination of EPA Risk Assessment Principles and Practices, US Environmental Protection Agency, Office of the Science Advisor, Washington, DC.

77. USEPA 2020 Technical overview of ecological risk assessment: risk characterization, United States Environmental Protection Agency, Washington DC.

78. USEPA, U. 1992. Guidelines for exposure assessment. Federal Register 57(104), 22888–22938.

79. Vos, T., Allen, C., Arora, M., et al. 2016. Global, regional, and national incidence, prevalence, and years lived with disability for 310 diseases and injuries, 1990–2015: a systematic analysis for the Global Burden of Disease Study 2015. The Lancet 388(10053), 1545-1602.

80. Wood, C.L., Sokolow, S.H., Jones, I.J., et al. 2019. Precision mapping of snail habitat provides a powerful indicator of human schistosomiasis transmission. P. Natl. Acad. Sci. USA 116(46), 23182–23191.

81. Xi-lin, D., Wei-ling, Z., Wei-dong, W., et al. 1999. Effects of temperature and dissolved oxygen content on oxygen consumption rate of Chinese prawn, giant tiger prawn and giant freshwater prawn. Chinese Journal of Oceanology and Limnology 17(2), 119–124.

82. Zhang, W. 2018. Global pesticide use: Profile, trend, cost/benefit and more. Proc. Int. Acad. Ecol. Environ. Sci. 8, 1–27.

83. Zhang, W.J., Jiang, F.B. and Ou, J.F. 2011. Global pesticide consumption and pollution: with China as a focus. Proceedings of the International Academy of Ecology and Environmental Sciences 1(2), 125–144.

