## Supplementary material for "Identifying low risk insecticides that can enhance food production without increasing mortality of biocontrol agents for human schistosomiasis": Table S1-S6

**Highlight:** Ecotoxicity of insecticides for biocontrol agents of the infectious human disease schistosomiasis integrates water and human systems via agricultural policies.

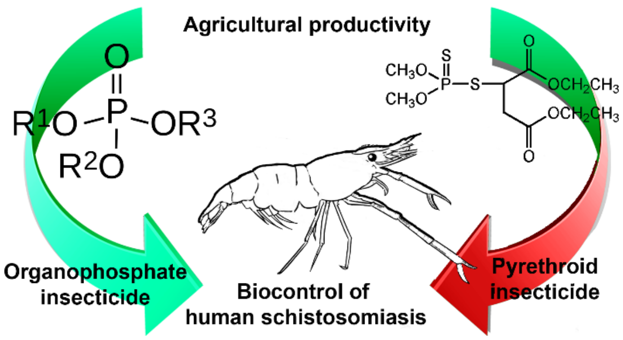

**Supplementary Information**

**Table S1.** Parameters for and results of calculations of annual peak estimated environmental concentrations (EECs) in ponds for each insecticide using the United States Environmental Protection Agency (US EPA) Pesticide in Water Calculator (PWC). All pesticide properties were obtained from the University of Hertfordshire Pesticide Property Database (2018), except as noted. The US EPA PWC calculates 30 years of estimated environmental concentrations using region-specific weather and soil data. We used standardized US EPA scenarios for growing corn from five states (Illinois, Mississippi, North Carolina, Ohio, and Pennsylvania), producing a total of 150 annual peak EEC estimates. Application rates were determined from manufacturer specimen labels from recommendations for application to corn.

| | Organophosphates | | | Pyrethroids | | |
|---|---|---|---|---|---|---|
| | Chlorpyrifos | Malathion | Terbufos | Esfenvalerate | λ-cyhalothrin | Permethrin |
| Sorption coefficient ($k_{oc}$; mL/g) | 8151 | 1800 | 500 | 251717 | 283707 | 100000 |
| Water column metabolism half-life (days) | 36.5 | 0.4 | -- | 56 | 15.1 | 40 |
| Water reference temperature (°C) | 20 | 20 | -- | 20 | 20 | 20 |
| Aqueous Photolysis Half-life (days) | 29.6 | 98 | 4.5 | 2 | 40 | 1 |
| Photolysis Reference Latitude (°) | 40 | 40 | 40 | 40 | 40 | 40 |
| Hydrolysis Half-life (days) | 25.5 | 6.2 | 6.5 | 428 | 0 | 31 |
| Surface Soil Half-life (days) | 76 | 3[a] | 5 | 66.6 | 175 | 13 |
| Foliar Half-life (days) | 10[b] | 1.5[b] | N/A | Unknown | Unknown | 8[c] |
| Soil Reference Temperature (°C) | 20 | 20 | 20 | 20 | 20 | 20 |
| Molecular mass (g/mol) | 350.59 | 330.4 | 288.45 | 419.9 | 449.85 | 391.28 |
| Vapor Pressure (torr) | 1.07E-05 | 2.30E-05 | 2.60E-04 | 8.78E-15 | 1.50E-09 | 5.25E-08 |
| Solubility (mg/L) | 1.05 | 148 | 4.5 | 0.001 | 0.005 | 0.2 |
| Henry's Constant (atm·$m^3$/mol) | 2.93E-06 | 4.89E-09 | 2.40E-05 | 4.10E-07 | 1.40E-05 | 1.40E-06 |
| Air Diffusion Coefficient ($cm^2$/day) | 1910 | 1296 | 3610 | 2520 | | |
| Heat of Henry (J/mol) | 52291.8 | 54041 | 54041 | 45727 | 49884 | 49884 |

Applications:

| Trade name (for determining application rates): | Lorsban Advanced | Cheminova Malathion 57 | Counter 15G | Asana XL | Warrior II w/ Zeon Tech. | Ambush |
|---|---|---|---|---|---|---|
| Number of applications | 3 | 2 | 1 | 5 | 16 | 8 |
| Days between applications | 10 | 5 | N/A | 5 | 4 | 5 |
| Amount (kg active ingredient/ha) | 1.12 | 1.12 | 1.46 | 0.06 | 0.03 | 0.28 |
| Application method | Foliar | Foliar | Incorporate | Foliar | Foliar | Foliar |
| Depth (cm) | N/A | N/A | 3 | N/A | N/A | N/A |
| Peak EEC (μg/L): | | | | | | |
| Maximum | 13 | 8.28 | 0.779 | 0.041 | 0.034 | 0.432 |
| Mean | 5.026 | 1.148 | 0.778 | 0.038 | 0.012 | 0.135 |
| Median | 4.615 | 0.518 | 0.778 | 0.038 | 0.012 | 0.109 |
| Minimum | 0.649 | 0.506 | 0.764 | 0.035 | 0.002 | 0.021 |

[a] – source: http://www.pesticideinfo.org/

[b] – source: http://pmep.cce.cornell.edu/profiles/extoxnet

[c] – source: Florida Department of Agriculture and Consumer Services

**Table S2.** Tested concentrations of each insecticide by class

| Pesticide | Class | Test Concentrations (µg/L) | | | | |
|---|---|---|---|---|---|---|
| Esfenvalerate | Pyrethroid | 0.02 | 0.20 | 1.00 | 2.00 | 20.00 |
| λ-cyhalothrin | Pyrethroid | 0.03 | 0.13 | 0.26 | 1.30 | 5.00 |
| Permethrin | Pyrethroid | 0.04 | 0.21 | 0.41 | 2.07 | 4.41 |
| Chlorpyrifos | Organophosphate | 0.64 | 3.20 | 6.40 | 32.00 | 64.00 |
| Malathion | Organophosphate | 10.10 | 101.00 | 1010.00 | 10100.00 | 101000.00 |
| Terbufos | Organophosphate | 0.09 | 0.86 | 8.55 | 17.10 | 171.00 |

**Table S3.** Summary of reported 96-h $LC_{50}$ values from USEPA ECOTOX Release 4.0 database for *Macrobrachium* spp. exposed to five insecticides.

| Chemical | Species | Exposure Type | $LC_{50}$ (ppb) | No. of studies |
|---|---|---|---|---|
| λ-cyhalothrin | *M. nipponense* | Renewed | 0.05 | 1 |
| Permethrin | *M. rosenbergii* | Static | 0.031 | 1 |
| Chlorpyrifos | None | | | 0 |
| Malathion | *M. lamarrei* | Static | 695.5 | 2 |
| Malathion | *M. ferreirai* | Static | 398 | 1 |
| Terbufos | None | | | 0 |

**Table S4.** Artificial spring water used to house *M. rosenbergii* throughout the experiment was composed of the indicated chemicals and concentrations in deionized water.

| Chemical | Concentration (ml / l) |
|---|---|
| Calcium Chloride | 1 |
| Magnesium Sulfate | 1 |
| Potassium Phosphate | 1 |
| Sodium Nitrate | 1 |
| Sodium Bicarbonate | 2 |
| Disodium metasilicate | 1 |
| Boric Acid | 1 |
| Potassium Chloride | 1 |

**Table S5.** Variance components from the mixed-effects Cox survival analysis for the laboratory study of *Macrobrachium rosenbergii* exposed to multiple concentrations of three pyrethroid (esfenvalerate, λ-cyhalothrin, and permethrin) and three organophosphate (chlorpyrifos, malathion, and terbufos) insecticides. Given below the random effects components are the coefficients for the fixed effect of concentration (toxic units) and the random effects for the intercepts of each class and chemical. Positive coefficients indicate a greater probability of mortality (relative to the grand mean) during the study per unit increase in toxic units. The magnitude of the difference in risk between two classes or chemicals can be found by taking the exponent of the difference of their respective coefficients.

| Parameter | Random effect | Standard deviation | Variance | Proportion of variance |
|---|---|---|---|---|
| Intercept | Chemical | 0.386 | 0.149 | 0.293 |
| Intercept | Class | 0.599 | 0.359 | 0.707 |
| Fixed and random effects | | coef | Hazard ratio | P-value |
| Fixed | Concentration | 0.594 | 1.811 | <0.001 |
| Class | Pyrethroid | 0.505 | 1.657 | N/A |
| Class | Organophosphate | -0.505 | 0.604 | N/A |
| Chem | Esfenvalerate | 0.302 | 1.353 | N/A |
| Chem | λ-cyhalothrin | 0.065 | 1.067 | N/A |
| Chem | Permethrin | -0.158 | 0.854 | N/A |
| Chem | Chlorpyrifos | 0.207 | 1.230 | N/A |
| Chem | Malathion | -0.327 | 0.721 | N/A |
| Chem | Terbufos | -0.090 | 0.914 | N/A |

**Table S6** Average values and cox model results for prawn field study prior to model selection (Wald tests substituted for LR tests to provide robust variances).

| Experimental level | Predictor | *Average (1 SE)* | *coef* | *Robust SE* | *Hazard (95%CI)* | *z-value* | *p-value* |
|---|---|---|---|---|---|---|---|
| Village | Total organophosphate use (ha) | 32.3 (7.5) | 0.11 | 0.04 | 1.11 (1.03-1.20) | 2.66 | 0.008 |
| **Village** | **Total pyrethroid use (ha)** | 5.2 (1.3) | 0.01 | 0.01 | 1.01 (0.99-1.03) | 1.02 | 0.306 |
| **Prawn** | **Water point temperature (°C)** | 28.2 (0.5) | 0.27 | 0.11 | 1.31 (1.06-1.63) | 2.53 | 0.012 |
| **Prawn** | **Sex of the prawn (male)** | | 0.42 | 0.21 | 1.52 (1.00-2.31) | 1.97 | 0.049 |
| Prawn | Weight of the prawn (g) | 23.3 (0.6) | 0.00 | 0.01 | 0.10 (0.99-1.01) | -0.39 | 0.693 |
| Prawn | Number of prawn claws | 1.4 (0.1) | -0.07 | 0.09 | 0.94 (0.79-1.11) | -0.76 | 0.447 |
| Village | Average percent slope cage to nearest field | 4.1 (0.6) | -13.98 | 6.82 | 0.00 (0.00-0.54) | -2.05 | 0.040 |
| Village | Average distance nearest field to water (m) | 18.1 (2.2) | -0.06 | 0.03 | 0.95 (0.90-1.00) | -1.99 | 0.047 |
| Village | Average distance cage to nearest field (m) | 4.1 (0.6) | 0.00 | 0.00 | 0.10 (0.99-1.01) | -0.09 | 0.928 |
| **Village** | Average emergent vegetation in 100m | 11,150.6 (898.3) | 0.00 | 0.00 | 1.00 (0.36-1.99) | 2.20 | 0.028 |
| **Village** | Average dissolved oxygen (ppm) | 0.5 (0.5) | -0.17 | 0.44 | 0.84 (1.00-1.02) | -0.39 | 0.698 |
| Village | Average conductivity (µS/cm) | 180.1 (30.0) | 0.01 | 0.00 | 1.01 (1.00-1.02) | 2.96 | 0.003 |
| Village | Interaction Avg. emergent vegetation-by-Avg. dissolved oxygen | | 0.00 | 0.00 | 1.00 (1.00-1.00) | -0.87 | 0.383 |
